# NuSeT: A Deep Learning Tool for Reliably Separating and Analyzing Crowded Cells

**DOI:** 10.1101/749754

**Authors:** Linfeng Yang, Rajarshi. P. Ghosh, J. Matthew Franklin, Chenyu You, Jan T. Liphardt

## Abstract

Segmenting cell nuclei within microscopy images is a ubiquitous task in biological research and clinical applications. Unfortunately, segmenting low-contrast overlapping objects that may be tightly packed is a major bottleneck in standard deep learning-based models. We report a <u>Nu</u>clear <u>Se</u>gmentation <u>T</u>ool (NuSeT) based on deep learning that accurately segments nuclei across multiple types of fluorescence imaging data. Using a hybrid network consisting of U-Net and Region Proposal Networks (RPN), followed by a watershed step, we have achieved superior performance in detecting and delineating nuclear boundaries in 2D and 3D images of varying complexities. By using foreground normalization and additional training on synthetic images containing non-cellular artifacts, NuSeT improves nuclear detection and reduces false positives. NuSeT addresses common challenges in nuclear segmentation such as variability in nuclear signal and shape, limited training sample size, and sample preparation artifacts. Compared to other segmentation models, NuSeT consistently fares better in generating accurate segmentation masks and assigning boundaries for touching nuclei.

## Introduction

Quantitative single cell analysis can reveal novel molecular details of cellular processes relevant to basic research, drug discovery, and clinical diagnostics. For example, cell morphology and shape are reliable proxies for cellular health and cell-cycle stage, as well as indicating the state of disease-relevant cellular behaviors such as adhesion, contractility, and mobility.^1,2,3,4,5^ However, accurate segmentation of cellular features such as the size and shape of the nucleus remains challenging due to large variability in signal intensity and shape, and artifacts introduced during sample preparation.^6,7^ These challenges are exacerbated by cellular crowding, which juxtaposes cells and obscures their boundaries. Additionally, in many traditional segmentation methods^8^ parameters need to be iteratively adjusted for images varying even slightly in quality.^9^

Convolutional neural networks (CNN) have emerged as a robust alternative to traditional segmentation methods for segmenting cell nuclei.^10,11,12,13,14,15^ CNNs achieve their superior performance through new deep-learning models.^10,16,17,18^ CNNs’ applicability for high precision image segmentation was first demonstrated by a Fully Convolutional Network (FCN) for pixel-level segmentation.^10^ Additional FCN cell segmentation models have since been developed.^14,19,20^ These models established a basic pipeline for CNN-based nuclear segmentation and achieved significant improvements in segmenting different types of cells including bacteria and mammalian cells.^14,20^ However, in their original form, FCNs typically required large training datasets to achieve high levels of accuracy.^10^ This bottleneck was overcome in U-Net by introducing a U-shaped network that incorporates pooling layers and up-sampling layers.^15^ Additionally in U-Net, the network was guided to segment overlapping objects by introducing weight matrices at cell-boundaries. Several state-of-the-art nuclear segmentation models have since been developed using this architecture.^11,13,21^ These models are curated and evaluated on pixel-level accuracy, where each pixel is treated equally without being classified as foreground or background. However, for cell biology, the main goal is to make reliable statements about *cells* as a whole (e.g. the number of cells, their average size and shape, detection of rare/unusual cells) rather than focusing on image pixels. For such problems, the idea of instance segmentation provides a more effective solution, as the loss function incorporates a sense of the whole object and not just individual pixels. Mask-R-CNN^18^ was developed to implement instance segmentation by fusing Region Proposal Networks (RPN) with FCN. By applying FCN-based segmentation on the regions proposed by RPN, Mask-R-CNN has achieved good segmentation results in real-world image datasets. However, Mask-R-CNN’s performance remains to be validated for nuclear segmentation, particularly for images with high cell density. Mask-R-CNN also employs a fixed size bounding box across all images, which is a serious limitation for samples with variable-sized nuclei.^17, 18^ Additionally, at the pixel level, the segmentation task of Mask-R-CNN is performed by FCN, which is less accurate with small training datasets compared with U-Net.^15, 22^

To address these issues, we have developed a <u>Nu</u>clei <u>Se</u>gmentation <u>T</u>oolset (NuSeT), which integrates U-Net^15^ and a modified RPN (based on the implementation of previous works ^23, 24^) to accurately segment fluorescently labeled nuclei. In this integrated model, U-Net performs pixel-segmentation, while the modified RPN predicts unique bounding boxes for each image based on U-Net segmentations. The resulting output provides seeds for a watershed algorithm to segment touching nuclei. To minimize segmentation errors stemming from fluorescence signal variability and cell density variability in samples, we employed a novel normalization method that uses only foreground pixel intensities for image normalization. To increase the robustness and applicability of the model, we used training sets including samples with wide variations in imaging conditions, image dimensions, and non-cellular artifacts. Extensive qualitative and quantitative evaluation suggest that our segmentation pipeline significantly improves nuclei segmentation, especially in distinguishing overlapping boundaries, and is generalizable to multiple sample types.

## Results

### NuSeT is a robust nuclear segmentation tool

Tools for segmenting fluorescent nuclei need to address multiple features and limitations of biological images.^6,25^ Typical issues and limitations include:

1. Boundary assignment ambiguity: biological samples frequently have very high cell density with significant overlap between objects.
2. Signal intensity variation: Within one image, the signal can vary within each nucleus (e.g. due to different compaction states of the DNA in heterochromatin vs. euchromatin) and across nuclei (e.g. due to cell to cell differences in nuclear protein expression levels and differences in staining efficiency).
3. Non-cellular artifacts and contaminants: Fluorescence microscopy samples are often contaminated with auto-fluorescent cell debris as well as non-cellular artifacts.
4. Low signal to noise ratios (SNRs): Low SNRs typically result from lower expression levels of fluorescent targets and/or high background signal, such as sample auto fluorescence. (Supplementary Figure 1).

We used an end-to-end training approach that incorporates both U-Net and Region Proposal Network (RPN)^15,17^ to address these issues (Methods). In our approach, the training and inference step consists of running an input image in parallel in both U-Net and RPN. The final output of U-Net consists of two feature maps of the same shape as the input image, representing background and foreground pixel-assignment scores.^10^ The final foreground prediction is then computed from the maximum class score of each pixel. Although U-Net alone performs well on some microscopy datasets^22,26^, we incorporated RPN since it was originally designed to detect objects in images with high information content.^17^ We reasoned that the accurate performance of RPN in detecting objects can be leveraged to improve nuclear segmentation performance. To achieve robust separation of touching nuclei, we used RPN bounding boxes to determine nuclear centroids, which were then supplied as seeds to a watershed algorithm.^27,28^ To improve segmentation accuracy in images with large nuclear size variations, we modified the original RPN architecture to use bounding box dimensions based on average nuclear size for each image (Supplementary Figure 2). Instead of training U-Net and RPN separately, we merged the feature-extraction part of RPN with the down-sampling part of U-Net to avoid longer training time and more memory cost (Figure 1a).^10,15,17,18^ In this way, the instance detection insights of RPN are extracted from the original U-Net structure.

**Figure 1.**
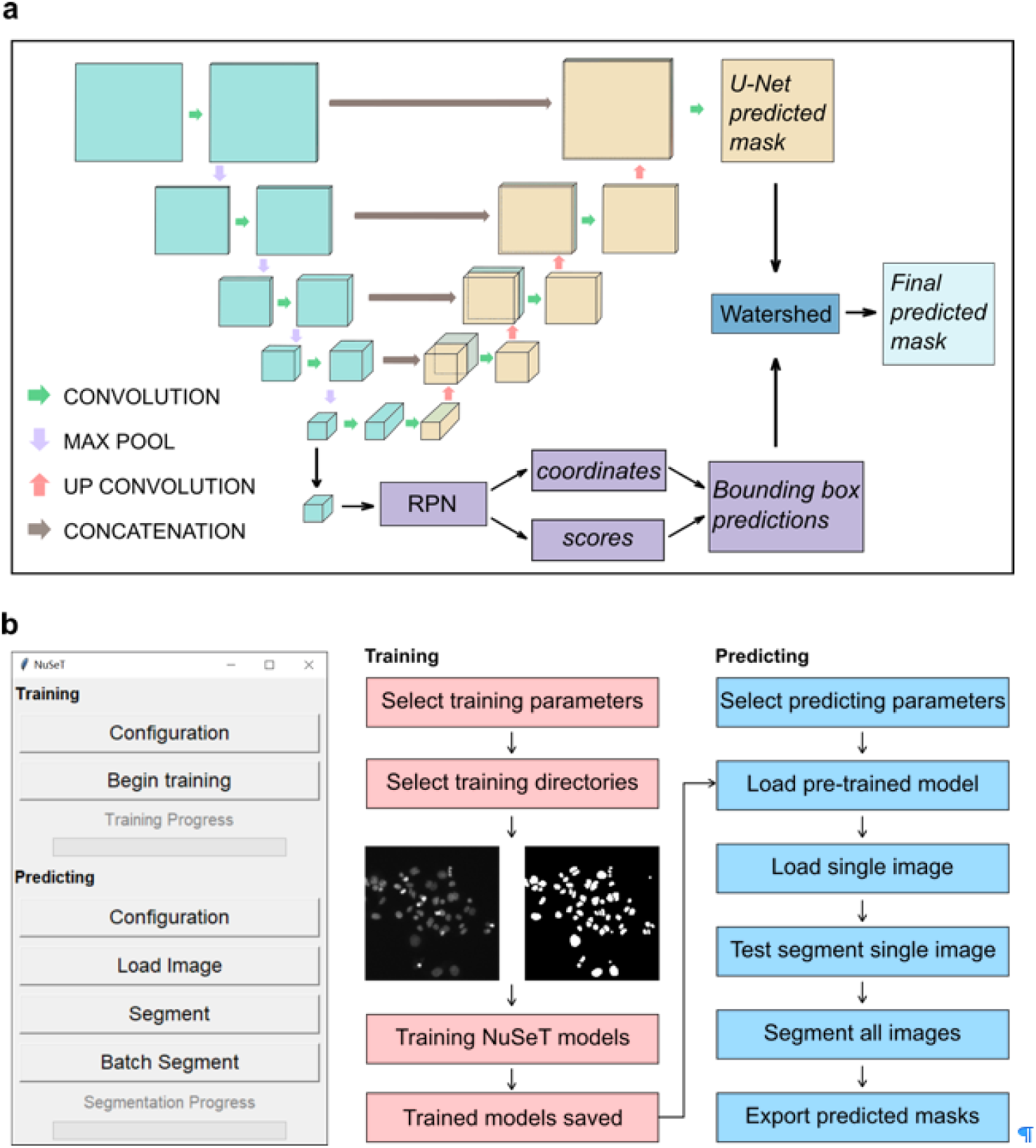
Pipeline for segmenting nuclei with NuSeT. **a**. Deep-learning model structure of NuSeT. The inputs of the model are gray scale images with different sizes. The outputs are binary masks with the same size as inputs, with predicted foreground regions as Ones and background regions as Zeroes. The model combines U-Net (gray and orange) and Region Proposal Network (purple), which performs nuclei segmentation and detection separately. The results are then merged and processed by watershed (dark blue) to generate final predictions. **b.** Outlook of NuSeT Graphic User Interface(GUI), and example training and predicting pipelines using NuSeT GUI.

**Figure 2.**
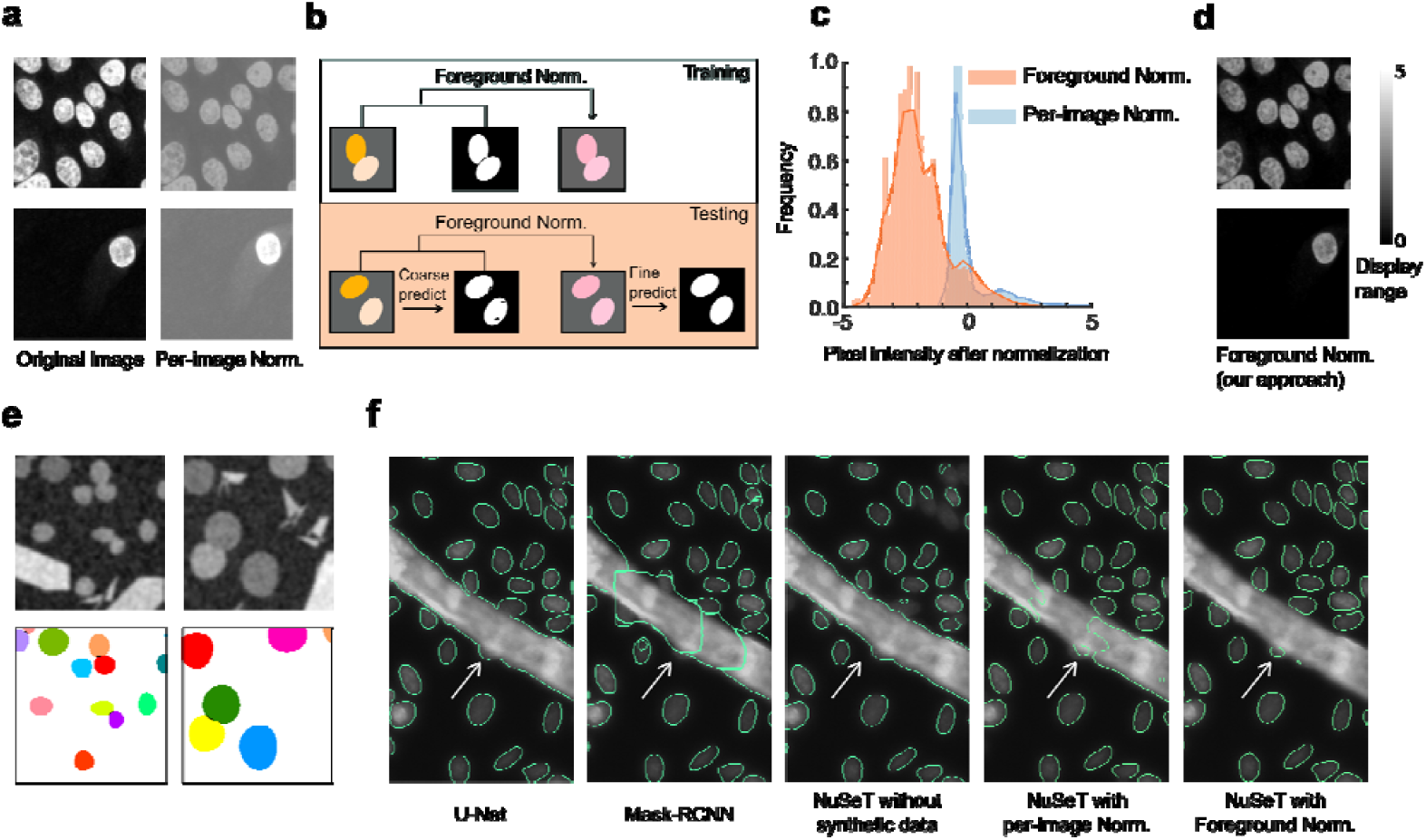
Improved normalization performance by foreground normalization and synthetic training. **a**. The visual effects of normalizing sparse/dense samples using Per-image normalization showing images having inconsistent nuclear signals after normalization. **b**. Foreground normalization during training and testing. During training, only pixels belonging to cell nuclei are used to normalize the image. During testing, a coarse segmentation prediction is generated by the model, and pixels belonging to the predicted nuclei are used to perform foreground normalization. The model then makes final predictions based on the normalized input images. **c**. Distribution of pixel intensities over an entire training dataset after different normalizations, showing foreground normalization has wider dynamic range. **d**. The visual effects of normalizing sparse/dense samples using foreground normalization showing images have higher dynamic range and more consistent nuclear signals. **e**. Examples of synthetic images with labels used during training. Our algorithm can generate synthetic nuclei-shaped blobs with different sizes, as well as different types of artifacts to increase the robustness of the model. Overlapping nuclei were introduced to enhance NuSeT performance in touching nuclei separation. **f**. Representative examples comparing the performances of different segmentation approaches. Training without synthetic images mis-identified artifacts (stripes) as foreground. The addition of synthetic data improved artifact detection. Switching to foreground normalization led the best performance including robust identification of imaging artifact, detected of more nuclei, and better separation of touching nuclei compared to Mask-RCNN and U-Net.

To evaluate the segmentation performance of the different algorithms, we computed the mean intersection over union for foreground and background (mean IoU), Root Mean Square Error (RMSE), and pixel accuracy (to benchmark pixel-level performance). Since in biological image processing most users (such as doctors and cell biology researchers) care much more about cell-level segmentation than pixel level accuracy, we also included instance-level segmentation metrics, such as the rate of correctly separating overlapping nuclei, and both the false-positive and false-negative detection rates (Methods). Two separate datasets, ‘MCF10A’ and ‘Kaggle’, were used to compare the performance of the algorithms. The MCF10A dataset consists of images of relatively uniformly fluorescent nuclei of a non-tumorigenic breast epithelial cell line^29^, grown to different levels of confluence. The Kaggle dataset was adapted from a public dataset^25^ representing cells from different organisms (including humans, mice, and flies) and containing images with a wide range of brightness, cell densities, and nuclear sizes. The overall comparison in Supplementary Table 1 and 2 suggests that NuSeT achieves similar segmentation accuracy compared with a current state-of-the-art pixel-level segmentation approach (U-Net), but achieves higher separation rates for overlapping nuclei. With the Kaggle dataset, NuSeT improved the separation of touching nuclei by more than 75% compared with U-Net. Compared with another state-of-the-art instance segmentation approach, Mask-RCNN, NuSeT achieved much lower false-negative detection rates in Kaggle dataset, leading to significantly better pixel-level segmentation accuracy. To make NuSeT more user-friendly, we have prepared a cross-platform graphic user interface (GUI) for the scientific community. Our GUI comes with the pretrained model which we used to benchmark NuSeT performance for various nuclei segmentation tasks. The GUI also allows the use of training and predicting modules (Figure 1b), allowing the users to perform custom segmentation tasks with NuSeT.

### Foreground normalization improves segmentation performance

Normalizing training data to alleviate image intensity differences is central to accelerating learning and improving network performance. Historically, imaging data have been normalized by subtracting the mean intensity calculated from all pixels in a dataset.^30, 31^ However, this leads to discrepancies in normalization, particularly for images with markedly different brightness levels. Normalizing data at per-image level addresses the issue of illumination differences^32^, but introduces brightness differences in images with sub-regions of varying cell densities (Figure 2a). Additionally, per-image normalization fares poorly in images strewn with auto-fluorescent artifacts (Supplementary Figure 3)

**Figure 3.**
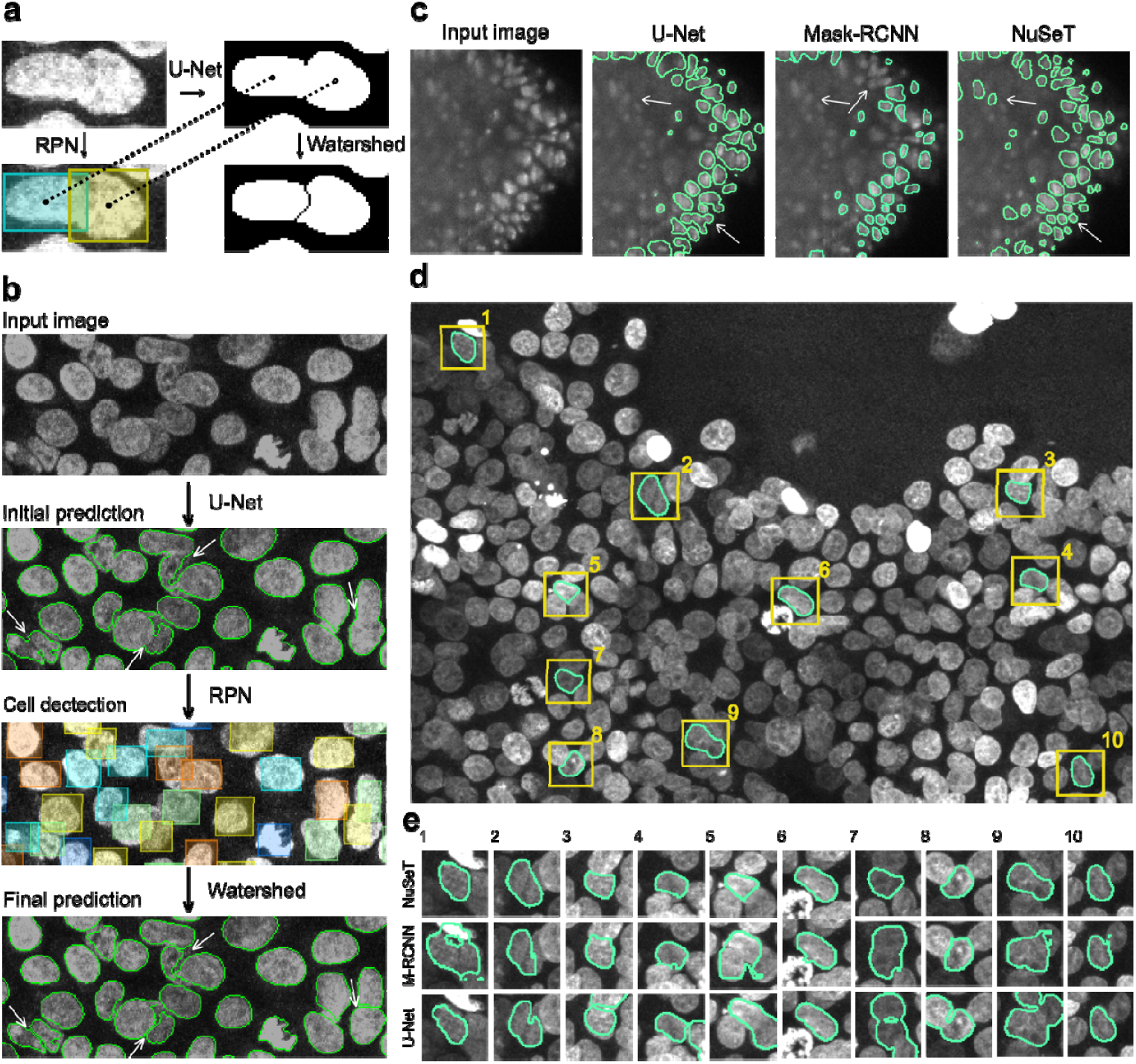
NuSeT efficiently addresses common segmentation challenges. **a.** Implementing RPN-aided watershed algorithm improves touching cell separation. Bounding boxes and segmentation mask are computed by RPN and U-Net. Then the estimated centroid of each cell is computed from the coordinates of the bounding box. The watershed line is then estimated based on the binary mask and centroids. **b.** Sample results showing that RPN successfully detects most of the cells, and watershed line further separate touching cells. **c.** Representative examples showing NuSeT detected more nuclei and better separated touching nuclei compared to Mask-RCNN and U-Net. **d, e.** Examples nuclear masks generated using NuSeT for an image with high nuclei density (d). Comparison with the corresponding masks generated by Mask-RCNN and U-Net show subtle as well as prominent irregularities in boundar delineation that are circumvented by NuSeT (e).

We incorporated a foreground normalization step in our data preprocessing. In this approach, only the pixels that belong to cell nuclei (foreground) are selected to calculate mean and standard deviation of pixel intensities. Since no label is provided during inference, foreground normalization requires two passes. In step one, the test data are normalized on a per image level to generate a coarse prediction of the foreground with our RPN-U-Net fusion. In step two, this coarse prediction is used to perform foreground normalization on test images before they are fed into the model for a second pass (Figure 2b). Compared with per-image normalization, the two-step foreground normalization approach is relatively robust to illumination differences, cell-density variations, and image artifacts and performs better in normalizing images with a broader dynamic range of pixel intensities (Figure 2c, d). As a result, model training with foreground normalization increased nuclei detection accuracy and boundary assignment for both Kaggle and MCF10A datasets (Supplementary Table 1). For datasets with high variations (notably, Kaggle), foreground normalization improved segmentation performance for all metrics considered, at both pixel and nuclei levels (Supplementary Table 1).

### Synthetic datasets in model-training improve detection and segmentation accuracy

Common sample contaminants have irregular shapes, significantly different overall brightness levels and aspect ratios compared to real cells, and uneven pixel intensities. To improve model performance and minimize false positive detection rates, we computationally generated synthetic images containing irregular shapes with varying intensities, as well as nuclei-like blobs (Methods). We also added gaussian blur and noise to the synthetic images to better represent real-world images. Additionally, overlapping blobs were included to mimic touching nuclei. Example synthetic images and training labels are shown in Figure 2e. Including synthetic data in the training process notably improved the model’s performance in distinguishing real nuclei from imaging artifacts (Figure 2f) and enhanced the separation of touching nuclei (Supplementary Table 1). The addition of foreground normalization on top of the synthetic images during model-training further reduced false positive detections (Figure 2f). Aided by these improvements, NuSeT outperformed both U-Net and Mask-RCNN in artifact detection/rejection (Figure 2f).

### RPN-aided Watershed improves boundary-resolution of highly overlapping objects

Having improved nuclear segmentation performance, we revisited the problem of separating overlapping nuclei. Recent work has used algorithms such as intervening and concave contour-based normalized cut^33, 34^ on binary segmentation masks extracted using traditional segmentation methods such as Otsu’s method^8^ to delineate overlapping nuclear boundaries. However, nuclear segmentation using traditional thresholding approaches failed to detect half of the nuclei in the Kaggle dataset (Supplementary Table 2), indicating that this approach is only effective for images with clean backgrounds and uniform signal.

We employed our modified RPN approach to detect nuclei. The nuclear centroids estimated from the RPN derived bounding box coordinates were passed as seeds for a watershed algorithm to separate touching nuclei.^27, 28^ Based on this, we added RPN-aided watershed as a post-processing step, which takes the U-Net segmented binary masks and RPN-predicted nuclei centroids to estimate the boundaries between touching nuclei (Figure 3a).

Our results suggest that a modified RPN can detect most nuclei in overlapping regions, and that a RPN-aided watershed separates 72%/94% of overlapping nuclei for Kaggle/ MCF10A dataset (Figure 3b, Supplementary Table 2). Compared with the modified RPN model without watershed, RPN-aided-watershed improved the overlapping nuclei separation performance by another 10% to 30% (Supplementary Table 2).

Through the integration of synthetic images, foreground normalization, and RPN-aided watershed, NuSeT consistently outperforms other state-of-the-art segmentation methods including U-Net and Mask-RCNN in nuclear boundary demarcation, particularly for blurry, low SNR nuclei (Figure 3c, Supplementary Figure 4, Supplementary Table 2). Mask-RCNN and NuSeT perform comparably in relatively sparse and homogenous samples (Supplementary Table 2). However, NuSeT approximates ground-truth boundaries more closely than U-Net and Mask-RCNN in samples with high cell densities (Figure 3d, e).

**Figure 4.**
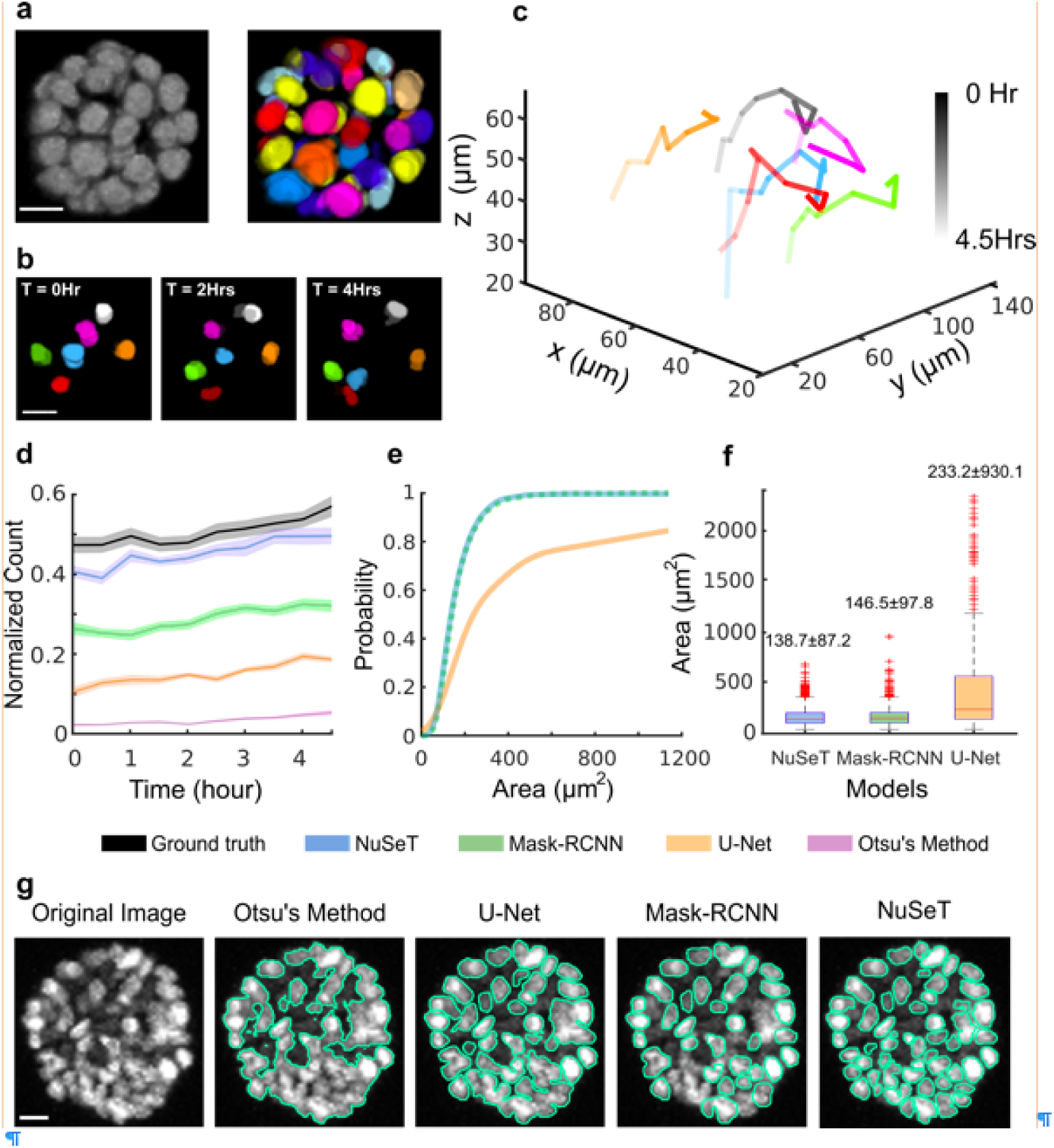
NuSeT effectively segments single nuclei in disorganizing dense mammary acini. **a.** Representative 3D MCF10AT acinus segmentation using NuSeT. **b.** Nuclei tracking. For ease of visualization only a few of the segmented nuclei are shown at different time points (See Supplementar Movie 2). **c.** 3D tracks of the nuclei shown in (**b**) over time, from 0 h (dark) to 4.5 h (light). **d.** Number of nuclei detected in disorganizing acini at different time points using different segmentation methods. Data were collected from 8 representative acini and were normalized by the total number of nuclei at the last time point. Data from the first 5 hours are shown. **e.** Cumulative distribution function plots of area of nuclei segmented using different methods. **f.** Box plots of nuclear area distribution. The median area for each method is indicated on the top. The area box plot for Otsu’s method (median area: 2816.6 ± 2845 μm^2^) is shown in Supplementary Figure 6. **g.** Representative examples comparing nuclei segmentation in dense mammary acini using different methods. Scale bars are 20 μm.

### Three-dimensional spatio-temporal tracking of individual nuclei in mammary acini

To investigate the performance of our algorithm in segmenting densely packed nuclei, we used NuSeT to segment and track nuclei in 3D reconstituted mammary acini grown from a Ras transformed MCF10A (MCF10AT) cell line. MCF10AT was chosen since upon continued growth in matrigel, this cell line produces mammary acini with very high cell density. Three-dimensional segmentation was performed by processing individual 2D slices from a Confocal Microscope Z-stack followed by three-dimensional reconstruction. NuSeT successfully segmented most of the nuclei in an acinus (Figure 4a), which facilitated seamless tracking of nuclei in mammary acini disorganizing on a 3D collagen matrix (Figure 4b, c). Both NuSeT and Mask-RCNN performed similarly on early stage mammary acini (cell count = ∼34 cells/acinus) (Supplementary Figure 5). To further evaluate the performance of different algorithms (NuSeT, UNET, Mask-RCNN and Otsu’s method) on segmenting nuclei in mammary acini, we carried out nuclear segmentation on 2D projections of dense mammary acini.

NuSeT accurately segmented most of the nuclei in dense mammary acini (Figure 4d-g). We were also able to track single nuclei through the entire process of acinar disorganization (*time* = 15 h, Supplementary figure 5, Supplementary movies 1, 2). Compared with the other widely used segmentation models, NuSeT performed consistently better at matching the number of nuclei detected manually for multiple acini (n = 8) at different stages of disorganization. U-Net, Otsu’s Method and Mask-RCNN all detected only a fraction of all the nuclei (Figure 4d) in a dense acinus. The distribution of areas of segmented nuclei (n = 1365 nuclei) across multiple acini (n = 8) at first 5 time points shows that while Mask-RCNN and NuSeT achieved comparable accuracy in nuclear boundary determination (median area of detected nuclei: 147 μm^2^ vs. 139 μm^2^, Figure 4e, f), Mask-RCNN only detected a subset of all nuclei (Figure 4d, g). Nuclear segmentation with U-Net on the other hand resulted in much larger nuclear area (median area of detected nuclei = 233 μm^2^, Figure 4e, f), indicating that U-Net often failed to separate touching nuclei (Figure 4g). All deep learning approaches outperformed the ‘traditional’ algorithm (Otsu’s Method, nuclei area = 2816.6 μm^2^, Supplementary Figure 5), as it rarely segmented single nuclei in dense settings (Figure 4g). Together, our results suggest that NuSeT outperforms both Mask-RCNN and U-Net in detecting nuclei and assigning boundaries for overlapping nuclei.

## Discussion

Here we present a deep learning model for nuclear segmentation that is robust to a wide range of image variabilities. Compared with previous models that need to be trained separately for specific cell types, NuSeT provides a more generalized approach for segmenting fluorescent nuclei varying in size, brightness and density. We have also developed novel training and pre/post-processing approaches to address common problems in biological image processing. Our results indicate that every stage in deep learning, from data collection to post-processing, is crucial to training an accurate and robust nuclear segmentation model. When compared with the state-of-the-art segmentation models, NuSeT separates touching nuclei better than U-Net and detects more nuclei than Mask-RCNN. Thus, it assimilates the advantages of both semantic segmentation (U-Net) and instance segmentation (Mask-RCNN) and circumvents their limitations. This combination enables NuSeT to analyze complex three-dimensional cell clusters such as mammary acini and track single nuclei in dynamic crowded environments. NuSeT will be easily adaptable to segment nuclei in other image types such as brightfield and H&E Stained samples using additional training data. We expect NuSeT to find wide applicability, particularly in the areas of cell lineage tracing and clinical diagnosis.

Although we have modified the original RPN architecture to adjust detection scales based on the median nucleus size for each image, NuSeT assumes similar nuclear sizes in the same image. This may account for the occasional errors in nuclei segmentation when using RPN-aided watershed. If markedly irregular (such as dim/deformed/blurry) nuclei are encountered in the same image, RPN may over- or under-detect the nuclei and produce incorrect numbers of bounding boxes. This would lead to marker misplacement and erroneous segmentation lines. While we expect NuSeT to perform well for nuclei of most mammalian cell types, its performance for mixed populations remain to be validated. Recent studies have extracted image features from multi-scale and ‘pyramidal hierarchy’ neural networks to improve detection accuracy for objects with large size variations.^35, 36^ Subsequent work has improved object detection in dense samples using weighted loss functions.^37^ By incorporating these advances into our current model, we expect to further improve NuSeT in multi-scale nuclei detection.

Our approach has cross-platform support and comparatively low hardware requirements. With a medium-level Nvidia GPU (Quadro P4000), training an accurate model only takes five hours, and the inference proceeds at 3 images/second. From a user’s standpoint, the NuSeT GUI enables researchers to easily segment their images without needing understand all the details of machine-learning, which connects state-of-the-art computer vision algorithms to a suite of cell biology problems. While in the present work we provide an effective and efficient pipeline for cell nuclei segmentation, this approach should be easily adaptable to a wide variety of image segmentation tasks involving densely packed and overlapping objects, such as jumbled piles of boxes or people in crowds.

## Methods

### Kaggle dataset preprocessing

The Kaggle dataset was downloaded from the Broad Bioimage Benchmark Collection (Accession number BBBC038v1).^25^ This dataset was sampled from a wide range of organisms include human, mice and flies, and the nuclei were recorded under different imaging conditions. Stage-1 training and test datasets were used for training and validation process. All the images were manually censored and training data with low segmentation accuracies were discarded. Only fluorescent images were used for training and validation process. We converted the run-length encoded labels to binary masks for both training and validation labels in MATLAB. The final Kaggle dataset used for our model contains 543 images for training and 53 images for validation. Segmentation errors, including mask misalignment and touching cells, were manually corrected image-by-image for training and validating data.

### MCF10A and mammary acini data collection

The data were collected on an Olympus FV10i confocal microscope with a 60X objective on MCF10A human breast epithelial cell line. The cell nuclei were stained with 1uM Sir-DNA for 1 hour before imaging. The test set consists of 25 experiments with the corresponding ground-truth binary labels. MCF10AT acini were grown and the acini disorganization assay were performed as described in Shi *et al*.^4^

### Data augmentation

To accelerate the training process, only simple data augmentation techniques were applied to the training images. We adopted mirror flip and small rotation (10 degrees, counterclockwise) for training data to alleviate the overfitting problem.

### Synthetic data generation

Synthetic cell nuclei images were generated by utilizing nuclei-like blobs (adapted from https://stackoverflow.com/questions/3587704/good-way-to-procedurally-generate-a-blob-graphic-in-2d), as well as random shape polygons/lines. Signal (brightness) variations were added to both blobs and polygons/lines. The sizes of nuclei like blobs, polygons and lines were varied image-by-image to simulate different imaging conditions. The synthetic images were generated with various image sizes, with width and height ranging from 256 pixels to 640 pixels. Gaussian noise and Gaussian blur were added to these images. We applied overlapping of blobs to strengthen the model capability in separating touching nuclei. The binary masks of the synthetic images were generated separately. To correctly separate all overlapping blobs in the corresponding segmentation masks, the positions of blobs were used as markers to apply watershed transform^28^ on overlapping blobs.

### Training and Inference details

To construct the training data, we incorporated 543 training images from Kaggle dataset and 25 training images from MCF10A dataset as the base-training dataset. After data augmentation, the training set contained 568 (original) + 568 (flip) + 568 (rotate) = 1704 images. Then we mixed the real images with synthetic images at 1:1 ratio to generate the final training dataset. The training images were normalized by subtracting the foreground mean value and dividing by the foreground standard deviation. Since U-Net contains 4 down-sampling and up-sampling layers, to make the tensors at each layer compatible, training images were further cropped so that widths and heights of the images were adjusted to the nearest multiple of 16. To train RPN, the ground truth coordinates for bounding boxes were calculated based on the binary nuclei masks. The coordinates of bounding box, (x_min, y_min), (x_max, y_max) were denoted as the most upper-left and lower-right pixels of the corresponding nuclei. Weight matrices were calculated per mask with w0 = 10 and sigma = 5 pixels as described in original U-Net paper.^15^ To avoid out-of-memory, one image was fed into the network at a time. During the training, the sequence order of the training data was reshuffled before each epoch to prevent overfitting. Learning rate was set to 5e-5, and Rmsprop^38^ was utilized as the training optimizer, and the best performance model was chosen within the first 30 epochs. The training loss was the sum of segmentation loss and detection loss. Segmentation loss was the sum of binary cross-entropy loss^10^ and Dice loss, and the detection loss was the class loss and regression loss as described in previous work.^17^

Two validation datasets were used to benchmark the model performance. The Kaggle validation dataset^25^ contains 50 images that have various type of nuclei under different imaging conditions. The MCF10A dataset contains 25 images that have homogenous nuclei imaged under the same setting manner. This study was performed on Nvidia Quadro P4000. Additional segmentation performance is shown in Supplementary Figure 6.

### Model evaluation

Eight models were chosen to compare their performance on both Kaggle and MCF10A validation dataset, including Otsu’s Method^8^, Deep cell^14^, U-Net^15^, Mask-RCNN^18^, NuSeT with per-image normalization and without synthetic data, NuSeT with per-image normalization, NuSeT with foreground normalization, and NuSeT with foreground normalization and RPN-aided watershed. The entire training dataset (with data augmentation and synthetic images) was applied to train all NuSeT models. To test Deep cell’s performance on the Kaggle and MCF10A dataset, we selected the model with highest pixel and nuclei level accuracy from http://www.deepcell.org/predict. Since no pre-trained two-dimensional fluorescent nuclei segmentation model was found from U-Net^15, 26^, we trained U-Net on our training dataset (without synthetic data) as our closest estimate for performance. The original Mask-RCNN model was trained for real-life segmentations. Therefore we trained Mask-RCNN on our training dataset (without synthetic data) starting from FPN-101 backbone.^39^

To evaluate model performance, we adopted the following performance metrics, including percentage of touching cell separated, false negative detection rate (F.N. rate), false positive detection rate (F.P. rate), mean I.U., RMSE, F1 and pixel accuracy. The first three metrics were evaluated on nuclei level, and the last four metrics indicate the performance on pixel level. The percentage of touching nuclei separated is calculated as:

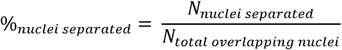

*N*_*nuclei separated*_ denotes the number of touching nuclei that have been successfully separated by the model, *N*_*total overlapping nuclei*_ denotes the total number of touching nuclei in the entire dataset.

F.N. rate is the proportion of the nuclei that the model fails to detect in the entire dataset. The detection failure is defined as: given a nucleus’ ground-truth binary mask, find the corresponding model-predicted mask that has the largest overlap ratio, which is measured by:

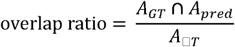

Where *A*_*GT*_ is the area of ground truth nucleus, *A*_*pred*_ is the area of model-predicted nucleus. If the overlap ratio is smaller than 0.7, it is suggested that the model fails to detect the nucleus. Hence the F.N. rate is denoted as:

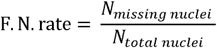

*N*_*missing nuclei*_ denotes the number of nuclei that the model fails to detect. *N*_*total nuclei*_ denotes the total number of nuclei labelled by ground-truth in the dataset.

Likewise, F.P. rate is the proportion of the nuclei that the model mis-detects in the entire dataset. The mis-detection is defined as: given a nucleus’ model-predicted mask, find the corresponding ground-truth mask that has the largest overlap, if the overlap ratio of the model predicted mask and the ground-truth mask is smaller than 0.7, it is suggested that the model detects an ‘nucleus’ that does not exist in the ground-truth. Hence the F.P. rate is denoted as:

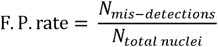

*N*_*mis-detections*_ denotes the number of model-predicted nuclei that found no match in the ground-truth labels.

Pixel-level metrics mean IU, F1, RMSE and pixel accuracy were calculated as:

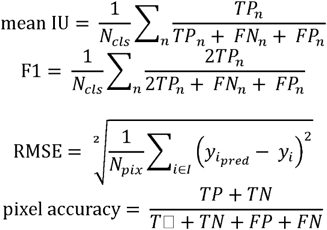

Where *TP, TN, FP, FN* denotes the pixel-level counts of true positive, true negative, false positive and false negative for single image. *N*_*cls*_ denotes the number of classes a pixel can be predicted to, in our case *N*_*cls*_ = 2 (foreground and background), and *TP*_*n*_ denotes the true positive counts of class n. *N*_*pix*_ is the number of pixels in the image, and 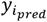 is the binary value of pixel i in the model-predicted mask, *y*_*i*_ is the binary value of pixel i in the ground-truth mask. The pixel-level metrics over the entire dataset were then calculated as the average metrics of all the images in the dataset.

## Code Availability Statement

All the code and pre-trained models will be released on github with detailed instructions so that researchers can easily apply them in practice upon publication.

## Acknowledgments

This work was partially supported by the National Institutes of Health (NIH) National Institute of General Medical Sciences (NIGMS)/National Cancer Institute (NCI) Grant GM77856, NCI Physical Sciences Oncology Center Grant U54CA143836, National Institute of Biomedical Imaging and Bioengineering (NIBIB)/4D Nucleome Roadmap Initiative 1U01EB021237.

## Author contributions

L.Y, R.P.G and J.T.L conceived the project. L.Y, R.P.G, J.M.F and C.Y designed research. L.Y performed most of the analysis with guidance from R.P.G. R.P.G collected 3D acini disorganization data. R.P.G and J.M.F collected MCF10A monolayer data. R.P.G, L.Y and J.T.L wrote the paper. All authors helped with editing.

The authors declare no conflict of interest.

## Supplementary Information

**Supplementary Table 1.** Internal performance comparison across different datasets.

| Per-image Norm. | Synthetic data | Foreground Norm. | Watershed | % of overlapping cells separated | FN rate | FP rate | Mean IoU | RMSE | F1 | Pixel accuracy |
| --- | --- | --- | --- | --- | --- | --- | --- | --- | --- | --- |
| √ |  |  |  | 37.90% | 15.12% | 7.28% | 0.88 | 0.15 | 0.94 | 0.97 |
| √ | √ |  |  | 51.49% | 17.04% | 8.56% | 0.88 | 0.16 | 0.93 | 0.96 |
|  | √ | √ |  | 64.13% | 13.96% | 8.37% | 0.89 | 0.15 | 0.94 | 0.97 |
|  | √ | √ | √ | 72.23% | 14.37% | 9.16% | 0.88 | 0.16 | 0.93 | 0.97 |

MCF10A dataset
| Per-image Norm. | Synthetic data | Foreground Norm. | Watershed | % of overlapping cells separated | FN rate | FP rate | Mean IoU | RMSE | F1 | Pixel accuracy |
| --- | --- | --- | --- | --- | --- | --- | --- | --- | --- | --- |
| √ |  |  |  | 76.50% | 1.57% | 8.96% | 0.93 | 0.18 | 0.96 | 0.96 |
| √ | √ |  |  | 84.14% | 2.46% | 6.48% | 0.94 | 0.17 | 0.97 | 0.97 |
|  | √ | √ |  | 91.70% | 4.34% | 6.14% | 0.93 | 0.18 | 0.96 | 0.96 |
|  | √ | √ | √ | 94.00% | 4.68% | 6.14% | 0.93 | 0.19 | 0.96 | 0.96 |

Step-by-step addition of synthetic data, foreground normalization, and RPN-aided watershed result in better separation of touching nuclei and more balanced false positive and false negative rates. Notice that the pixel-level accuracies (mean IU, RMSE, F1, pixel accuracy) are similar, despite marked differences in correctly calling cell boundaries and derived metrics (e.g. cell numbers).

**Supplementary Table 2.** External performance comparison of published models across different datasets.

|  | % of overlapping cells separated | FN rate | FP rate | mean IU | RMSE | F1 | Pixel accuracy |
| --- | --- | --- | --- | --- | --- | --- | --- |
| Otsu's Method | 14.18% | 50.81% | 15.27% | 0.79 | 0.22 | 0.86 | 0.93 |
| Deep-cell | 24.11% | 4.47% | 30.51% | 0.79 | 0.21 | 0.87 | 0.94 |
| U-Net | 40.21% | 11.29% | 8.48% | 0.89 | 0.15 | 0.94 | 0.97 |
| Mask-RCNN | 80.04% | 23.9% | 4.54% | 0.85 | 0.17 | 0.92 | 0.96 |
| NuSeT | 64.13% | 13.96% | 8.37% | 0.89 | 0.15 | 0.94 | 0.97 |
| NuSeT+ watershed | 72.23% | 14.37% | 9.16% | 0.88 | 0.16 | 0.93 | 0.97 |

MCF10A dataset
|  | % of overlapping cells separated | FN rate | FP rate | mean IU | RMSE | F1 | Pixel accuracy |
| --- | --- | --- | --- | --- | --- | --- | --- |
| Otsu's Method | 41.74% | 21.10% | 6.65% | 0.83 | 0.29 | 0.90 | 0.91 |
| Deep-cell | 89.15% | 9.16% | 5.37% | 0.85 | 0.28 | 0.92 | 0.92 |
| U-Net | 84.96% | 2.00% | 7.91% | 0.94 | 0.17 | 0.97 | 0.97 |
| Mask-RCNN | 95.65% | 2.94% | 5.51% | 0.91 | 0.21 | 0.95 | 0.95 |
| NuSeT | 91.70% | 4.34% | 6.14% | 0.93 | 0.18 | 0.96 | 0.96 |
| NuSeT+ watershed | 94.00% | 4.68% | 6.14% | 0.93 | 0.19 | 0.96 | 0.96 |

### Supplementary Figures

**Supplementary Figure 1.**
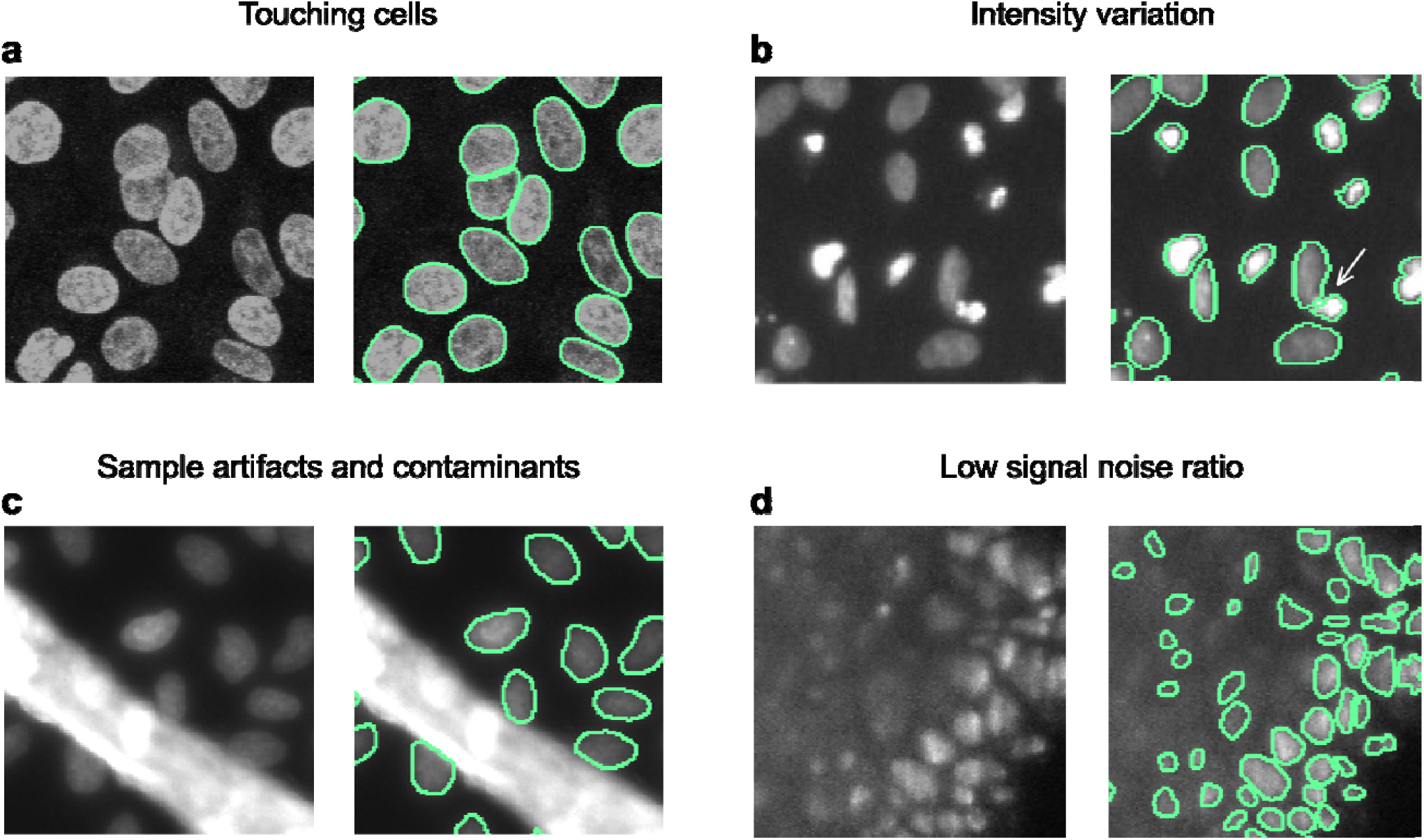
Common problems encountered in nuclei segmentation. Some common factors that affect the quality of nuclei segmentation, are, touching cells (**a**), signal variation (**b**), sample preparation artifacts and contaminants (**c**), and low signal to noise ratio (**d**). Colored outlines represent the goals (ground truth) for segmentation tasks.

**Supplementary Figure 2.**
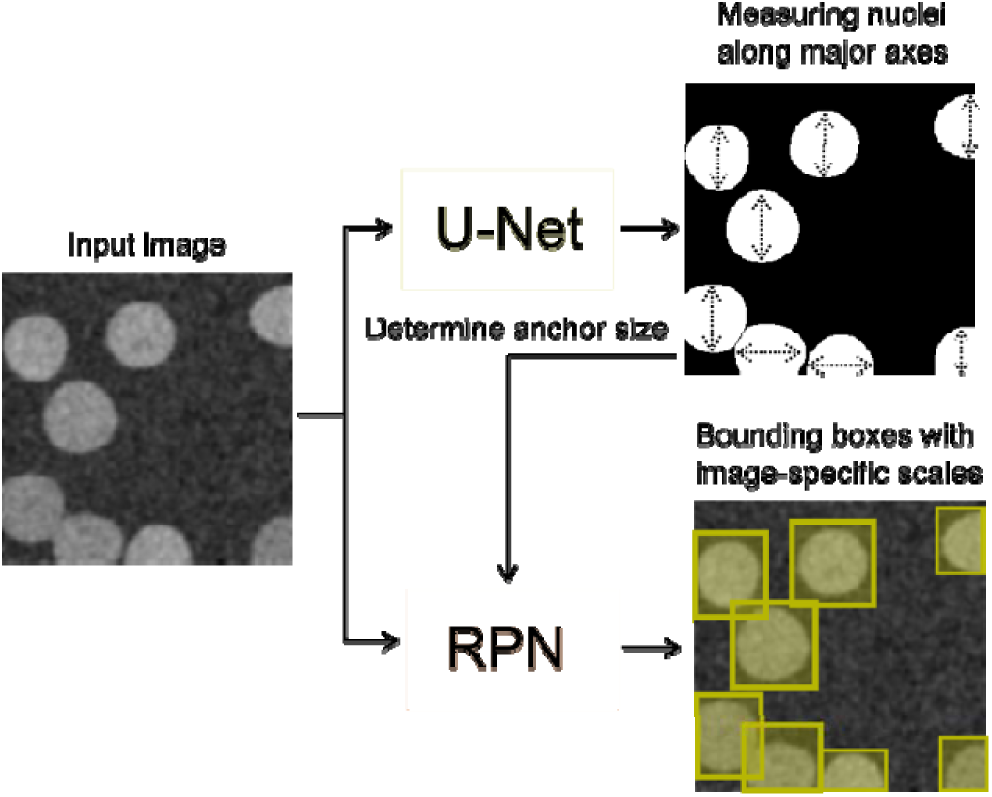
Adjusting bounding box dimensions based on nuclear size. Historically RPN has used a rigid base size for all bounding boxes, which resulted in high detection error rate in the Kaggle dataset. We improved the RPN so that it applies different bounding box base sizes for different images. The base size is determined by the median of all nuclei sizes within the image. Nuclei sizes are defined by the maximum value between nuclei widths and heights.

**Supplementary Figure 3.**
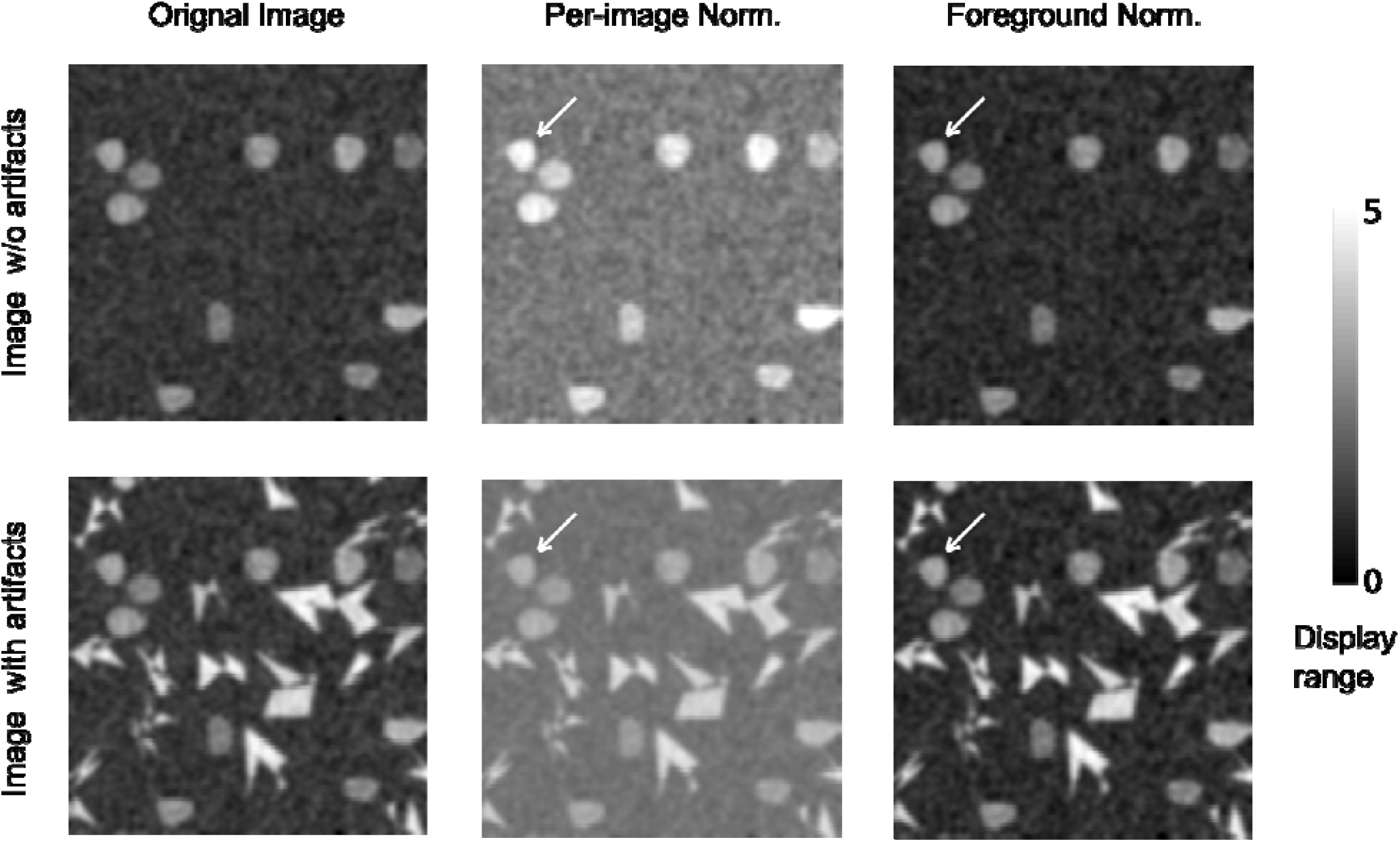
Foreground normalization is more robust than per-image normalization in handling images with sample preparation artifacts. Normalizing samples with or without sample artifacts using different normalization methods show that images have more consistent nuclei signals after foreground normalization (highlighted by arrows).

**Supplementary Figure 4.**
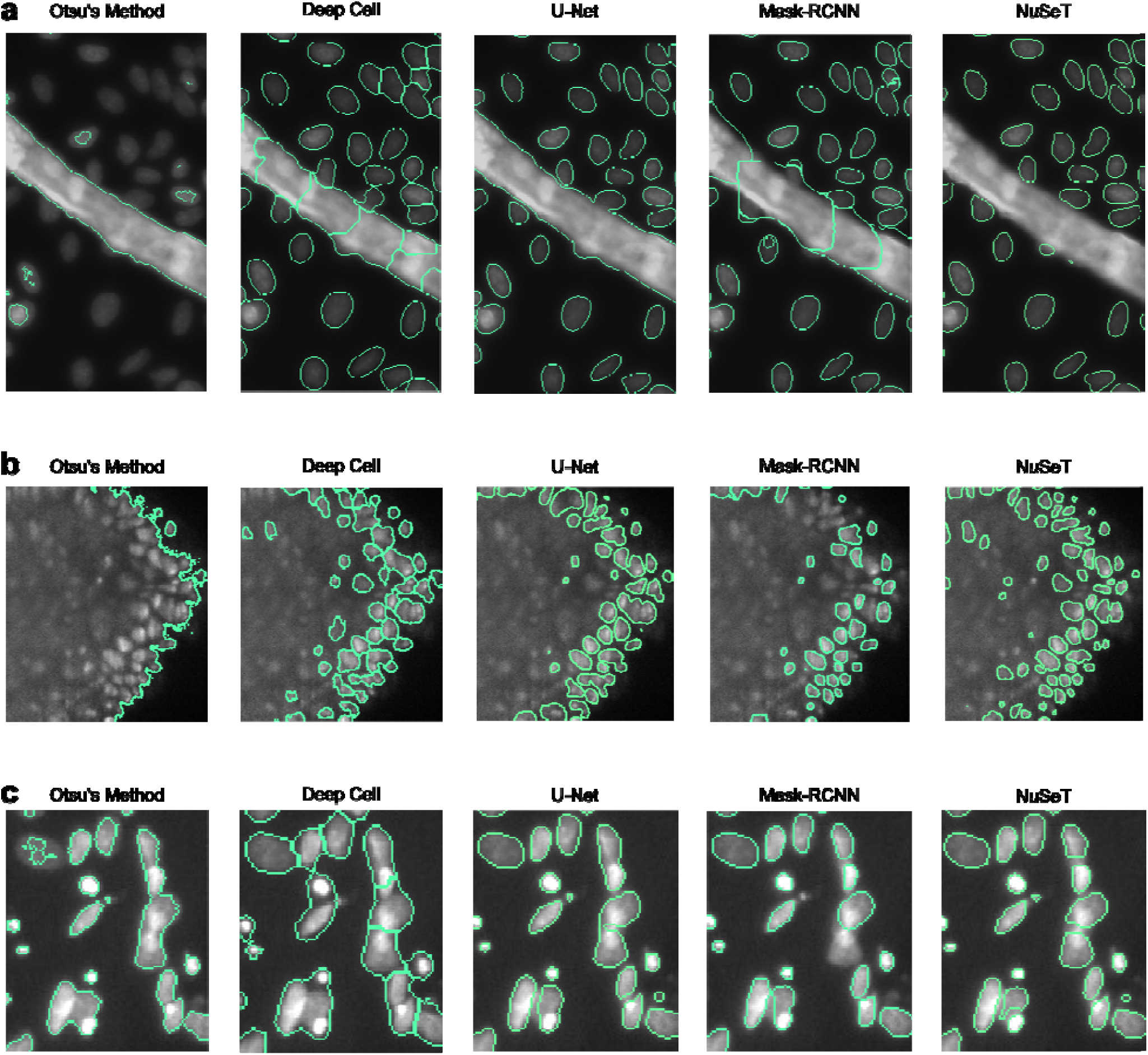
Additional segmentation performance comparisons across algorithms, including traditional thresholding approach (Otsu’s method) and Deep cell.

**Supplementary Figure 5.**
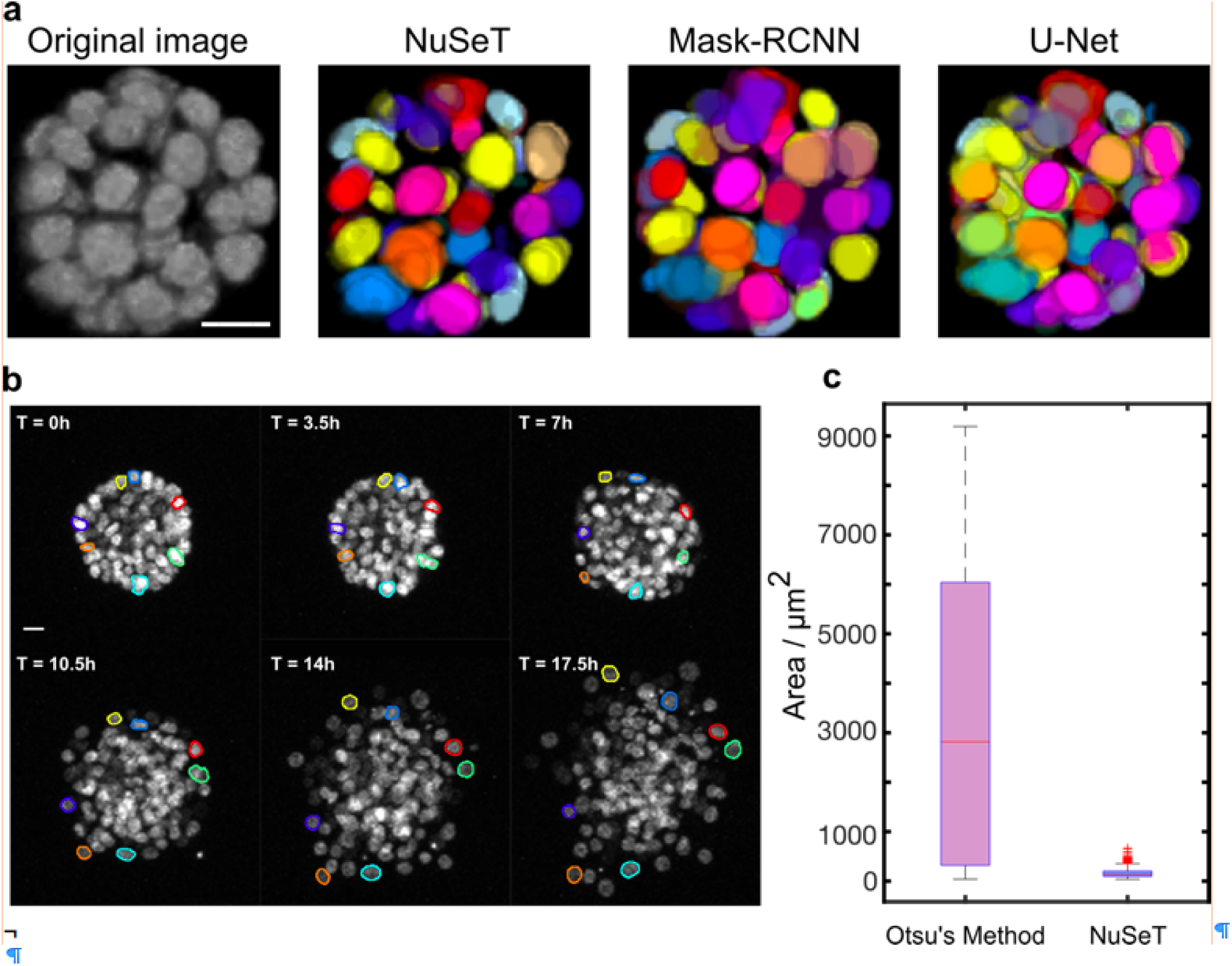
Additional mammary acini segmentation and tracking results. **a.** Three-dimensional acini tracking with different deep-learning models. **b.** Additional time-lapse tracking of selected nuclei. **c.** Comparison of nuclei area distribution for Otsu’s method (median area: 2816.6 ± 2845.0 μm^2^) and NuSeT (median area: 138.7 ± 87.2 μm^2^).

**Supplementary Figure 6.**
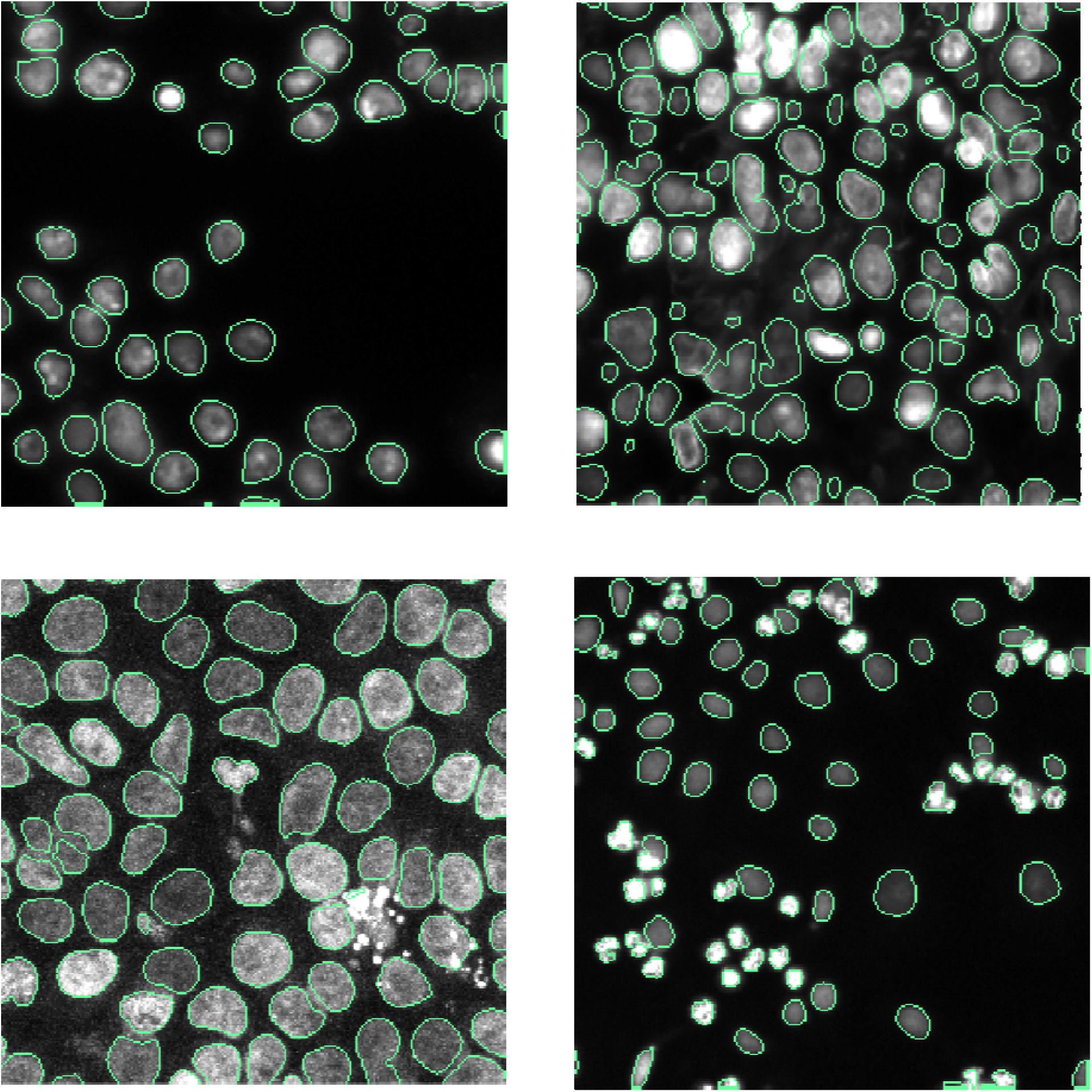
Additional examples showing NuSeT’s performance when handling images with signal variations, shape variations, touching nuclei and sample preparation artifacts.

**Supplementary Movie 1:** Time-lapse tracking of sample nuclei segmented using NuSeT in a 2-D maximum projection of a disorganizing mammary acinus.

**Supplementary Movie 2:** 3-D Time-lapse tracking of sample nuclei segmented using NuSeT in a disorganizing mammary acinus.

### Supplementary notes about the NuSeT user interface (UI)

**Figure 1.**
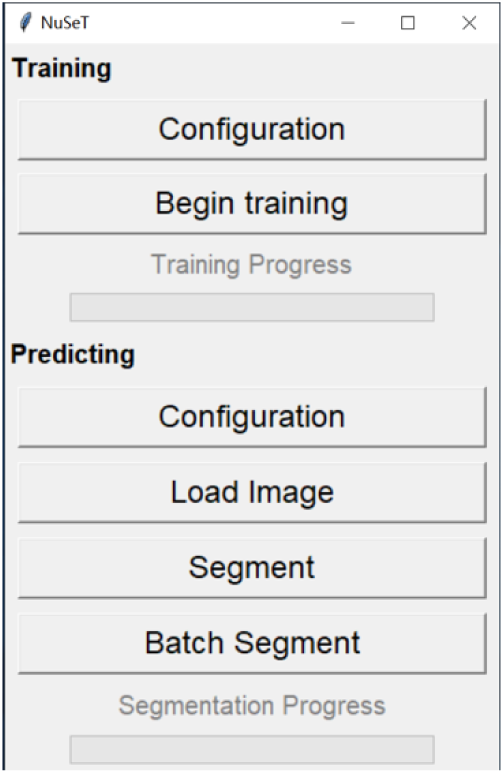
The main menu of NuSeT software. Once all the software dependencies are met, move to folder NuSeT/, and issue the command ‘*python3 NuSeT.py*’ in the command line. This User Interface (UI) has been separately tested for Ubuntu (Linux), Mac and Windows, and passed all tests in three platforms. If Windows 10 is used, the NuSeT UI will pop up as in Figure 1. There are two modules in the UI, namely **Training** and **Predicting**. The training module has two functions: **Configuration** allows you to choose basic training parameters, including number of training cycles (epochs), learning rates, and optimizers (Figure 2). Currently only Adam and Rmsprop optimizers are allowed, since they proved to work better in practice. Click the **Save** button when finished setting training parameters. Alternatively, user can choose the default parameters (35 epochs, learning rate 0.0001, Rmsprop) by simply ignoring this function.

**Figure 2.**
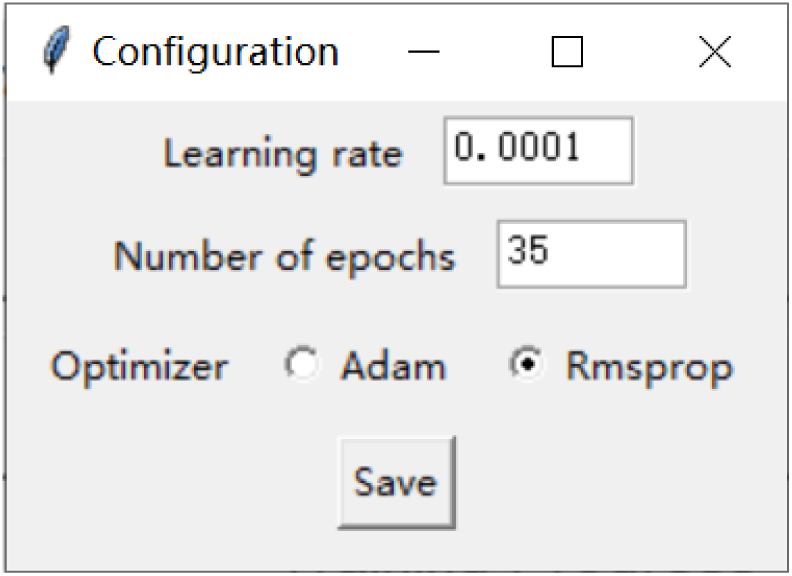
The configuration page for NuSeT training module. Before the training, the training images and training labels should be stored in separate folders, and the name of the images and the corresponding training labels should be the same. After closing the configuration window, click **Begin training**, and NuSeT will ask for the training image directory and training label directory. Upon choosing both directories, training will automatically start (Figure 3). The neural network will start with training a whole image normalization model, and then apply the whole image normalization results to train the foreground normalization model. This training step can take many hours if user’s training dataset is big.

**Figure 3.**
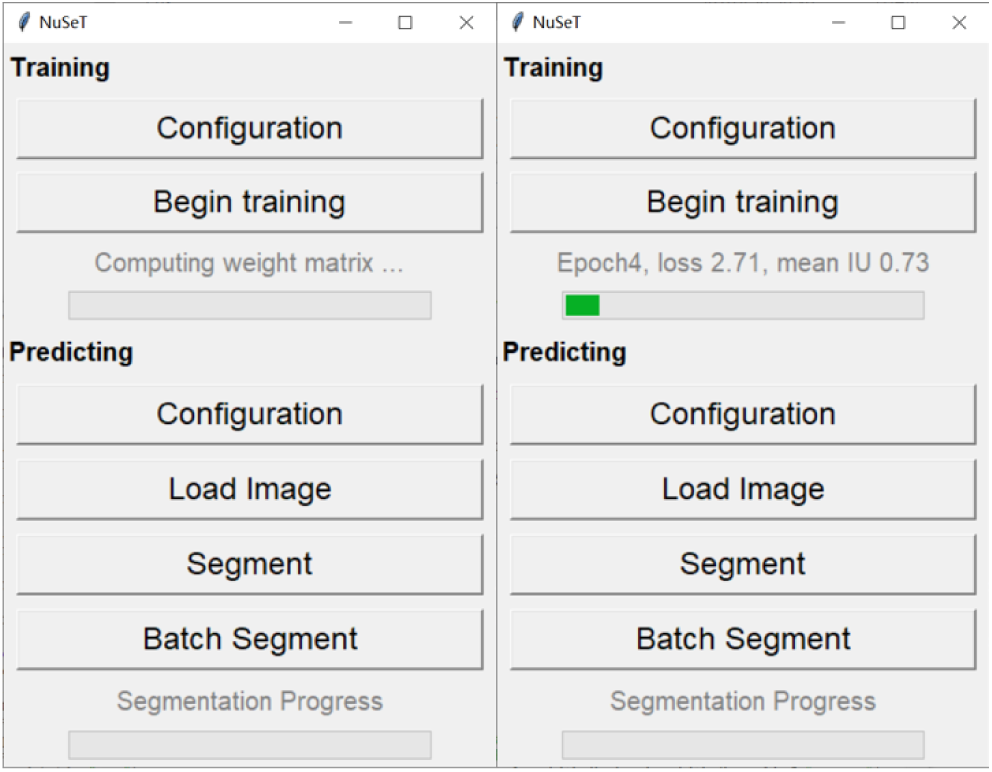
Sample UI displays during the training process. Once the training is finished, the trained model will be saved at Network/ folder, and then we are ready to move to the Predicting module. Predicting module also allows you to choose several segmentation parameters by clicking the configuration button (Figure 4). First it asks the user whether they need the RPN-aided watershed transform. Then NuSeT asks the user to define the following parameters: Min detection score and Non-max-suppression (NMS) ratio, which are used to control the cell-detection confidence. Briefly, lower min detection score and higher NMS ratio allows more cells to be detected and separated, but the segmentation error will also increase accordingly. Click save to finish the parameter setup. Likewise, users can leave the default parameters (with watershed, Min detection score 0.85, NMS overlapping ratio 0.1) by ignoring this function.

**Figure 4.**
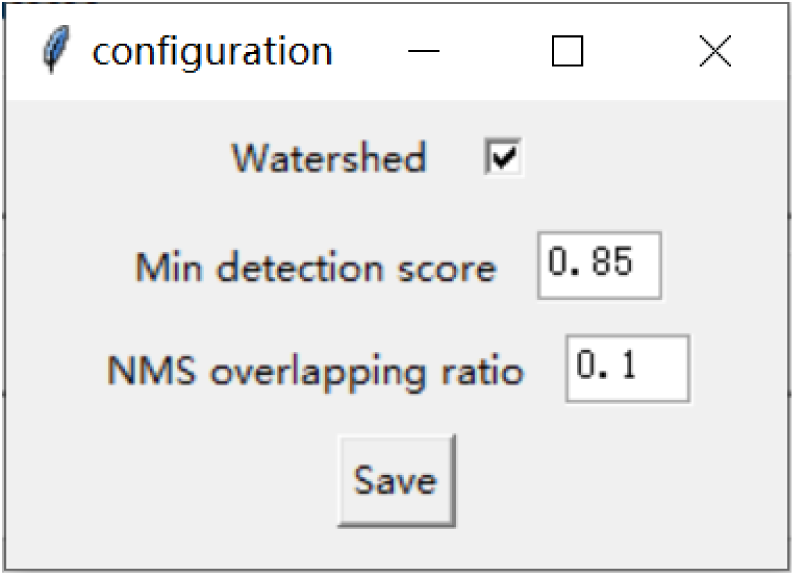
The configuration page for NuSeT predicting module. We have provided pretrained model that was used in the paper to segment fluorescent nuclei, which was saved in the default Network/ folder, however user can apply their model from the training module above, which will automatically overwrite the original model when training is finished. NuSeT also allows user to visualize sample segmentation results for single image by clicking the Load Image button and Segment button (Figure 5).

**Figure 5.**
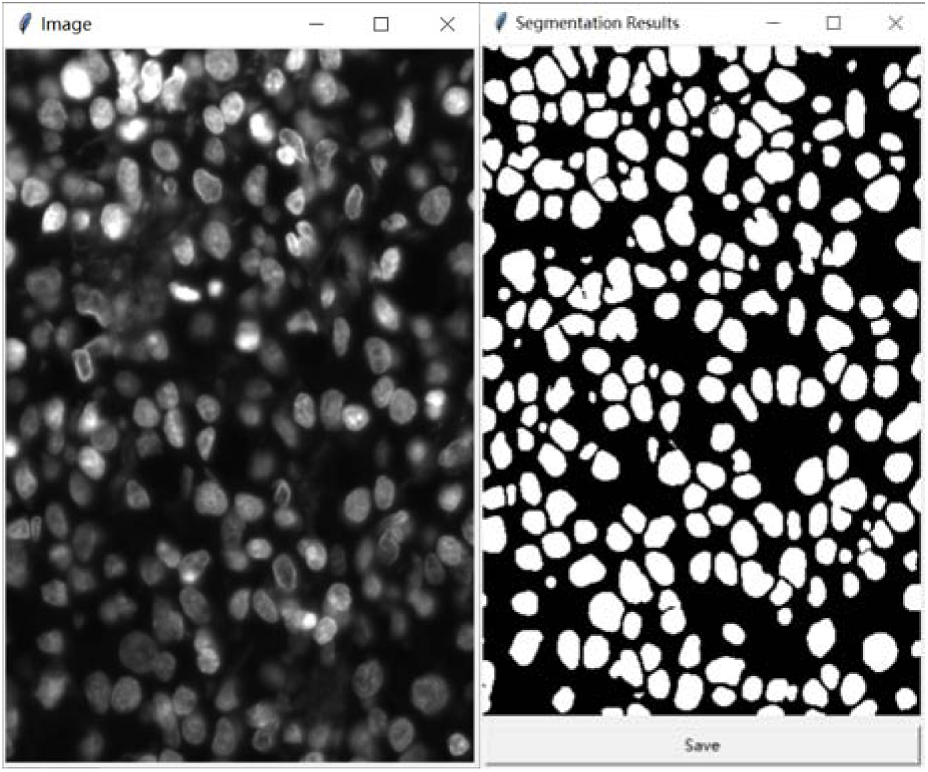
Example segmentation result using pre-trainined NuSeT model. Once the segmentation results have been validated, NuSeT allows the user to segment all images within the given directory (**Batch Segment** button), and all segmentation results (binary masks) will be stored in the same directory as the images.

